# State-dependent protein-lipid interactions of a pentameric ligand-gated ion channel in a neuronal membrane

**DOI:** 10.1101/2020.04.07.029603

**Authors:** Marc A. Dämgen, Philip C. Biggin

**Author notes:** Corresponding author; (PCB).

## Abstract

Pentameric ligand-gated ion channels (pLGICs) are receptor proteins that are sensitive to their membrane environment, but the mechanism for how lipids modulate function under physiological conditions in a state dependent manner is not known. The glycine receptor is a pLGIC whose structure has been resolved in different functional states. Using a realistic model of a neuronal membrane coupled with coarse-grained molecular dynamics simulations, we demonstrate that the lipid-protein interactions are dependent on the receptor state, suggesting that lipids may regulate the receptor’s conformational dynamics. Comparison with existing structural data confirms known lipid binding sites, but we also predict further protein-lipid interactions including a site at the communication interface between the extracellular and transmembrane domain. Moreover, in the active state, cholesterol can bind to the binding site of the positive allosteric modulator ivermectin. These protein-lipid interaction sites could in future be exploited for the rational design of lipid-like allosteric drugs.

**Author Summary:** Ion channels are proteins that control the flow of ions into the cell. The family of ion channels known as the pentameric ligand gated ion channels (pLGICS) open in response to the binding of a neurotransmitter, moving the channel from a resting state to an open state. The glycine receptor is a pLGIC whose structure has been resolved in different functional states. It is also known that the response of pLGICs can also be modified by different types of lipid found within the membrane itself but exactly how is unclear. Here, we used a realistic model of a neuronal membrane and performed molecular dynamics simulations to show various lipid-protein interactions that are dependent on the channel state. Our work also reveals previously unconsidered protein-lipid interactions at a key junction of the channel known to be critical for the transmission of the opening process. We also demonstrate that cholesterol interacts with the protein at a site already known to bind to another compound that modulates the channel, called ivermectin. The work should be useful for future drug design.

## Introduction

Members of the pentameric ligand-gated ion channel (pLGIC) family are pervasively found in all neurons and their primary function is to mediate neuronal communication (1, 2). They are located in the postsynaptic membrane, where neurotransmitter binding leads to conformational changes that open a channel pore permitting passive ion passage that leads to cell hyperpolarization or depolarization (3). Their dysfunction is associated with a wide range of neurological diseases such as epilepsy, Parkinson’s, Alzheimer’s and schizophrenia, and thus they constitute pivotal drug targets (4-6). In mammals, the pLGIC superfamily consists of the cation-selective, excitatory nicotinic acetylcholine receptors (nAChRs), 5-hydroxytryptamine type 3 receptors (5-HT_3_Rs) and a single zinc-activated channel as well as the anion-selective inhibitory γ-aminobutyric acid type A receptors (GABA_A_Rs) and glycine receptors (GlyRs). Moreover, the invertebrate glutamate-gated chloride channel (GluCl) from *Caenorhabditis elegans* as well as the prokaryotic ion channels from *Gloeobacter violaceus* (GLIC) and *Erwinia chrysanthemi* (ELIC) are other important pLGICs (7).

All pLGICs consist of five subunits that are arranged in a pseudosymmetric manner around the pore axis. They share a conserved overall architecture: a ligand-binding extracellular domain (ECD), a transmembrane domain (TMD) containing the ion channel pore and an intracellular domain (ICD), which is completely absent in prokaryotic pLGICs. For protein-lipid interactions with the membrane environment, the TMD is of particular interest. Each of the five subunits consists of four membrane spanning helices (M1-M4). These helices can be subdivided into 3 concentric rings with respect to their position in the membrane according to Barrantes (8). The inner ring consists of the M2 helices which line the pore through which ions flow. They are shielded from the membrane by the M1 and M3 helices that form the middle ring. The latter are partly in contact with the membrane, being shielded towards the inside by the M2 helices and towards the outside by the M4 helices. The M4 helices make up the outer ring and expose the largest surface area towards the membrane environment and are only shielded from the lipid environment towards the inside by contacts with the M1 and M3 helices from the middle ring.

PLGICs are modulated by their lipid environment, but exactly where lipids bind, and how they act, is poorly understood (9-11). Via reconstitution experiments, cholesterol (CHOL) and anionic lipids have been demonstrated to be crucial for nAChR function (12-15). In pure phosphatidylcholine (PC) or phosphatidylethanolamine (PE) membranes, nAChRs bind agonist, but remain in an unresponsive, non-conductive state (that has been termed the uncoupled state). However, ternary mixtures of PC, cholesterol and the anionic lipid phosphatidic acid (PA), completely restore nAChR functionality. Interestingly, binary mixtures of PC/CHOL and PC/PA alone are not sufficient to stabilize a large pool of agonist-responsive nAChRs. Furthermore, from biophysical studies the M4 transmembrane helix has emerged as playing a role as a lipid sensor (16, 17). Since mutations along this helix alter gating, M4-lipid interactions are likely to influence the gating pathway (18-20). Detailed mechanistic insight at a molecular level remains elusive, but recent experimental evidence supports the so-called M4-M1/M3 lipid sensor model which hypothesizes that enhancing M4-M1/M3 interactions potentiates pLGIC function (21). These interactions are facilitated by aromatic residues lining this interface (22), but if they are lacking, then lipids could potentially act in a surrogate manner to modulate the enhancement of M4-M1/M3 interactions. Experimental evidence for this hypothesis is provided by studies of GLIC and ELIC in pure PC membranes (21). ELIC, reconstituted in pure PC membranes, has very few aromatic residues at the M4-M1/M3 interface, just like the nAChR, and is equally unresponsive upon agonist binding. GLIC, on the other hand, has more aromatic residues at the M4-M1/M3 interface and, remains functional in pure PC membranes. However, by engineering additional aromatic residues at the M4-M1/M3 interface of ELIC, its functionality can be restored.

Increasingly more structural data of pLGICs with resolved lipids, neurosteroids and detergents is becoming available, providing insights into the molecular details of protein interactions with lipids or lipid-like molecules. Known binding sites for phospholipids are located at the extracellular half of the TMD on the surface of the M1 and M4 helices as seen in a recent GABA_A_R structure (PDB 6I53 (23)) or inserted between the M4 helix and the adjacent M1 and M3 helices in several GLIC structures (e.g. PDB 3EAM (24), 6HZW (25). Phospholipids can also bind at the interface of adjacent subunits at the extracellular TMD half between (+)M3 and (-)M1 as showcased by the crystal structure of GluCl (PDB 4TNW (26) and which overlaps with the binding site of the positive allosteric modulator ivermectin in GlyR (PDB 5VDH (27) and 3JAF (28)) and GluCl (PDB 3RIF (29)). Interestingly, the anionic lipid phosphatidylserine (PS) is a competitor for the ivermectin binding site in GluCl (26), illustrating the competitive nature for lipid occupation on the receptor surface. Moreover, the ivermectin binding site overlaps with the interaction site of neurosteroids. Cholesteryl hemisuccinate is a detergent that is often used to replace cholesterol in crystallization studies and is bound to the ivermectin binding site in a GABA_A_R chimera (PDB 5OSA (30)), suggesting that cholesterol itself might be able to interact at the ivermectin binding site. Furthermore, the endogenous neurosteroid allotetrahydrodeoxycorticosterone (THDOC) acts as a potent positive allosteric modulator of GABA_A_Rs and binds in a similar location. Anionic lipids are found in the intracellular membrane leaflet and accordingly, a phosphatidylinositol 4,5-biphosphate (PIP2) binding site was identified in the intracellular half of the TMD next to the M3 and M4 helices of the α1 subunits in the GABA_A_R structure (PDB 6I53 (23)). Two cholesterol molecules have recently been resolved at the intracellular half of theTMD near the M3 and M4 helices of a cryo-EM structure of the nAChR (PDB 6CNJ and 6CNK (31) as suggested previously (21). An extensive list of all current pLGIC structures with lipids resolved can be found in (10).

While insights from the increasing amount of structural data are valuable, most are obtained under non-physiological conditions. It is not clear for example whether the artificial crystal lattice organization in crystal structures accurately reflects positional preferences of lipids in a natural membrane environment. When *in vivo*, different lipid types compete over interaction sites on the protein surface, but experimental conditions rarely account for this. For example, when obtaining the crystals for GluCl (PDB 4TNW) only 1-palmitoyl-2-oleoyl-sn-glycero-3-phosphocholine (POPC) was added as a single lipid and bound to the ivermectin binding site (26), yet other lipid types might have a higher affinity to this site. Unnatural detergent molecules might preferentially bind to what would be actual lipid sites in a natural membrane environment. The most recent cryo-EM structures of receptors reconstituted in lipid nanodiscs provide the closest mimic to a natural membrane environment, yet the influence of artificial surface tension on protein structure and dynamics is yet to be determined. It is rare for lipid densities from experimental techniques to allow for unambiguous determination of headgroups or chain length and thus, assignment of the actual lipid type. It is not uncommon that lipid density is simply modelled as a PC molecule with no justification (see for example the supplementary information of (24) or the methods section of (23)). Moreover, lipid interactions are of a highly dynamic nature (32), which poses a limit on the resolution that can be obtained for lipid densities.

Molecular dynamics (MD) simulations provide a complementary tool to experimental techniques that can probe protein-lipid interactions in a natural membrane environment under physiological temperature and pressure (33). In particular, coarse-grained (CG) MD simulations (using, e.g. the popular Martini model (34)) allow for longer time scales to be explored, which is required to rigorously sample the protein-lipid interaction space (32). They have been successfully applied to provide structural insights into protein-lipid interactions of key functional membrane proteins, such as transporters (35), ion channels (36) and G-protein coupled receptors (GPCRs) (37) to name a few.

While it is experimentally known that pLGICs are sensitive to their lipid environment and that cholesterol and anionic lipids are both essential for nAChR function, it is still the standard approach to perform simulations of this receptor class in simple bilayer models that only consist of a single lipid class, usually POPC (38, 39). Few simulations have included the functionally crucial lipid types, commonly in very simplified membrane models of binary or tertiary composition with no asymmetry between leaflets (40-42). Yet, *in vivo*, plasma membranes consist of many different lipid types that are distributed asymmetrically across the leaflets. However, so far no simulations of pLGICs in such a realistic membrane environment have been performed. Based on extensive lipidomics data, Ingollfson et al. (43) have derived a coarse-grained model of a neuronal membrane with asymmetric leaflet composition which can be used as an in vivo-mimetic model for the natural environment of pLGICs.

Here, we have used molecular simulation to reconstitute the GlyR, in an *in vivo*-mimetic neuronal membrane (see **Fig. 1**) and performed CG-MD simulations to probe the protein-lipid interactions. Our membrane model is a simplified version of (43) that consists of representatives for the 5 main lipid types found in neuronal plasma membranes: PC, PE, sphingomyelin (SM), PS, and CHOL. The reduction of the model to these 5 lipid types is necessary to ensure sufficient sampling of the protein-lipid interactions for each lipid type. Yet, it still contains all essential lipid types, particularly the functionally relevant cholesterol and anionic PS, in a typical asymmetric composition of a neuronal membrane. Moreover, to examine possible state-dependent protein-lipid interactions, we have performed two separate sets of simulations, each containing a different state of a human homology model of the α1 GlyR. One is derived from the inactive (closed) state (PDB 5CFB (44)) and one is derived from the active (open) state (PDB 3JAE (28)). Lipid densities and local residence times reveal that the different conformational states recruit a different lipid environment, thus supporting the view that lipids play a modulatory role in stabilizing different receptor states. We report previously unreported phospholipid interactions at the ECD-TMD interface that are likely to play a role in stabilizing the tertiary structure in this critical region for signal transduction. Furthermore, cholesterol can bind to the binding site of the positive allosteric modulator ivermectin and thus might itself act as a membrane-intrinsic modulation molecule. Noteworthy is also that the anionic lipid PS can bind near the intracellular ends of the M3 and M4 helices which overlaps with the PIP2 binding site in human GABA_A_R, but the details of the headgroup interactions differ. By elucidating a detailed map of the state-dependent protein-lipid interactions with a pLGIC, we provide insights that could be exploited for rational drug development of lipid-like receptor modulators.

**Figure 1.**
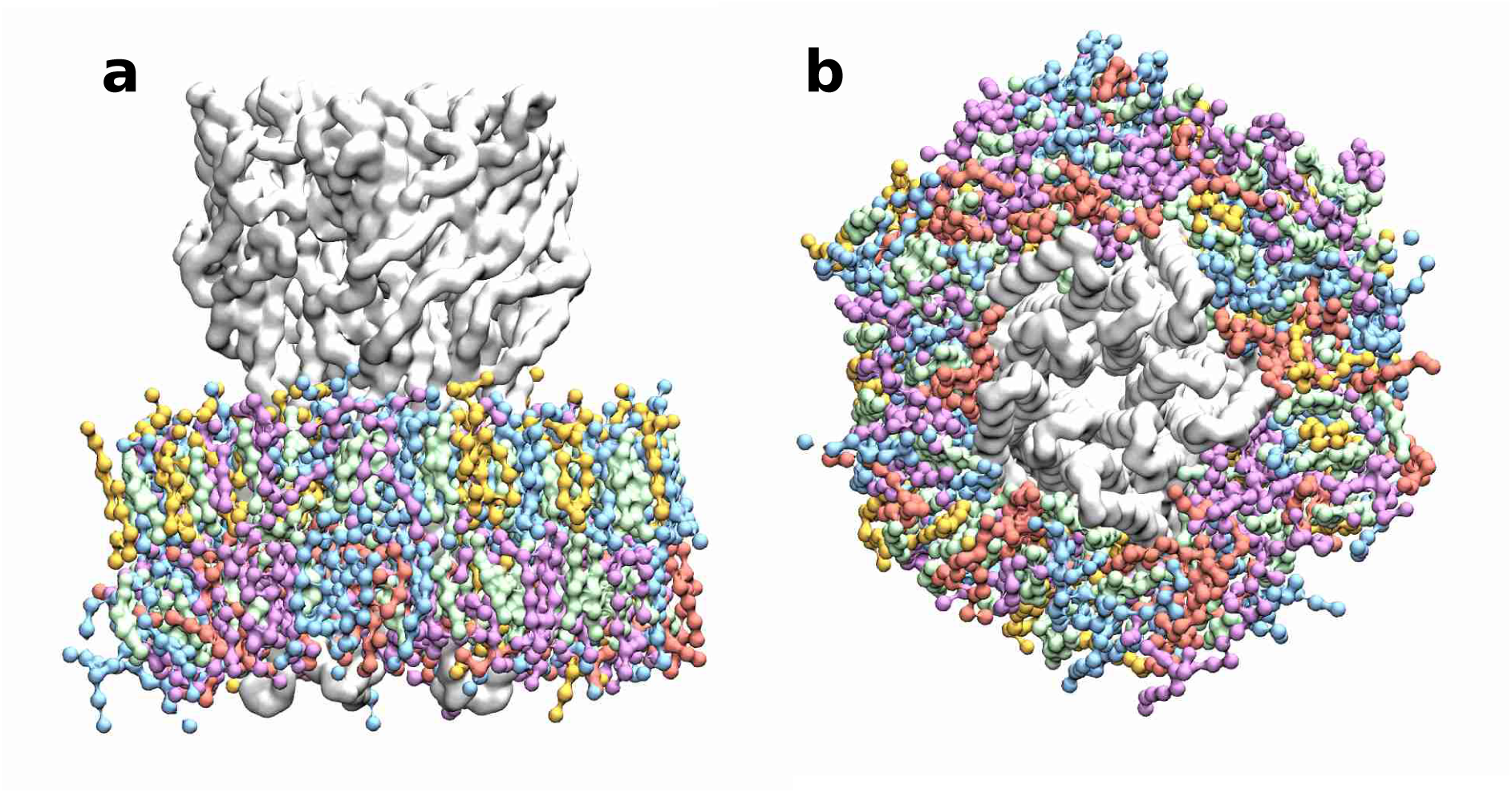
Simulation System. The simulation system **a** viewed in parallel to the membrane plane and **b** viewed from the intracellular side. GlyR (light grey) is embedded in a neuronal model membrane (PC: blue, PE: purple, SM: gold, CHOL: green, PS: red), water and ions are not shown for clarity.

## Results

### Bound lipids show dependence on the receptor state

To explore the modulatory role of lipids on GlyR receptor activation, we analysed the distribution of each lipid type per leaflet for both the inactive and active state (**Fig. 2**). Hot spots that are strongly preferred by certain lipid types can be clearly identified. Although the location of interaction hot spots does not vary much for most of the lipids (PC, PE, SM and PS), the relative occupation probabilities at these locations are clearly dependent on the state of the receptor, indicating that the binding affinities of these locations are sensitive to the conformational state of the receptor. We observe that PC, PE and SM can all occupy the same interaction hot spots, indicating they compete for the same binding locations. The fact that these 3 lipid types engage in similar interactions with the receptor is not surprising, given the similarity of their headgroups: PC and SM both have a phosphocholine headgroup, while PE has a phosphoethanolamine headgroup, which is chemically very similar. Note that in the coarse-grained Martini force field employed here, these groups only differ by single bead (particle type), namely ‘Q0’ for the choline group in PC and ‘Qd’ for the ammonium group in PE

**Figure 2.**
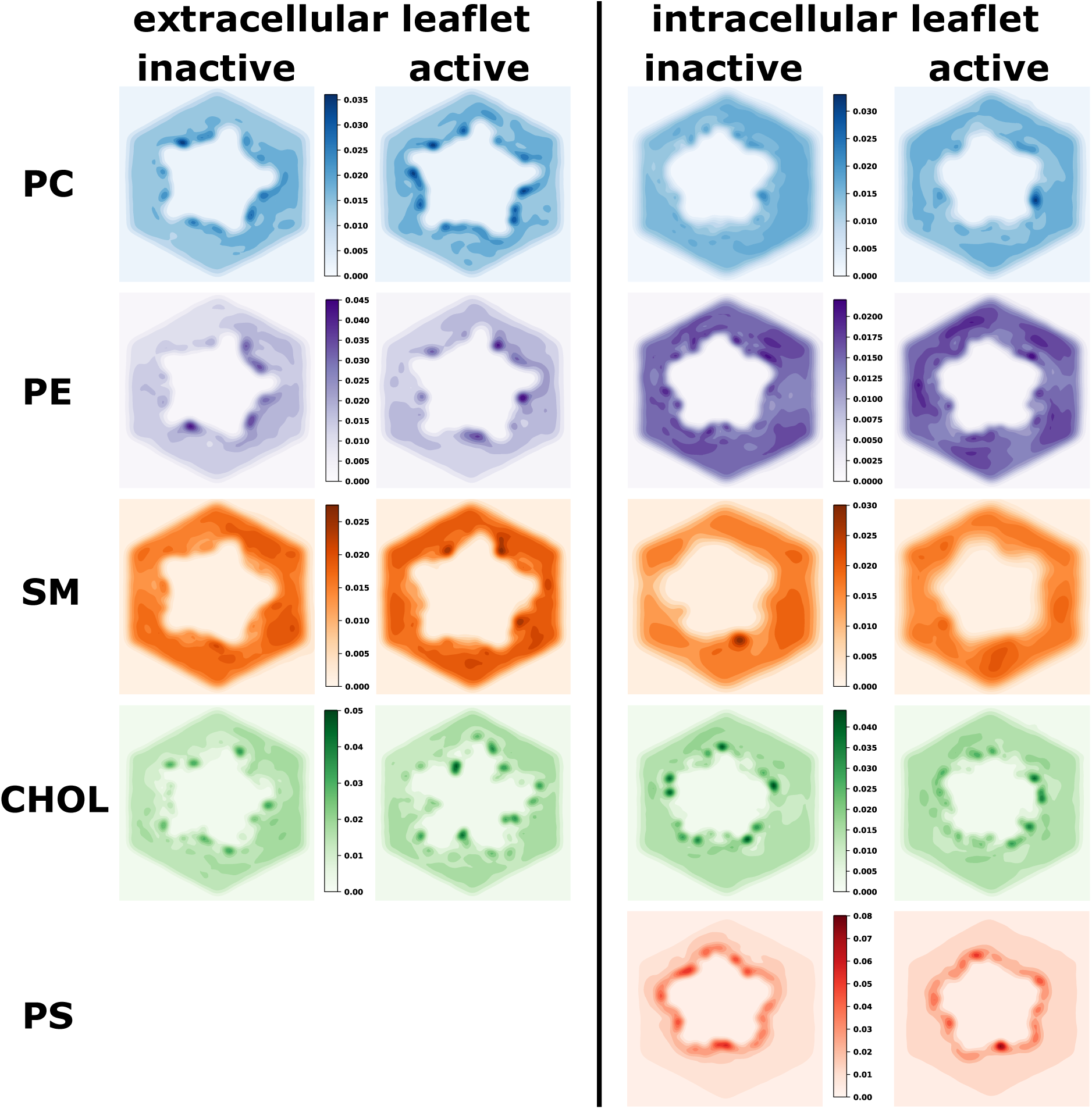
Probability densities of different lipid types around the receptor in the extracellular and intracellular leaflet for the inactive and active state. The colour scales were chosen to highlight the state-dependent differences per leaflet and lipid type. Darker colours represent interaction hotspots. For cholesterol, the location and intensity of interaction hot spots changes between the inactive and active state. PC, PE and SM can occupy similar sites on the receptor surface, but their intensity varies between states, suggesting a state-dependent affinity change.

If simulations were perfectly converged, we would expect to observe C5-symmetry around the 5-fold symmetric receptor. While this is achieved quite well for CHOL, this is not quite the case for the other lipid types, which are much less represented in the overall membrane composition and thus likely require longer simulation times to fully converge. This is despite using 10 × 40 μs of simulation time per receptor state and that well exceeds typically reported timescales of CG-MD simulations to study protein-lipid interactions, which are typically in the range of 5-10 repeats of 2-10 μs (37, 42). Nevertheless, some striking differences can be observed (**Fig. 2** and **Fig. 3**).

**Figure 3.**
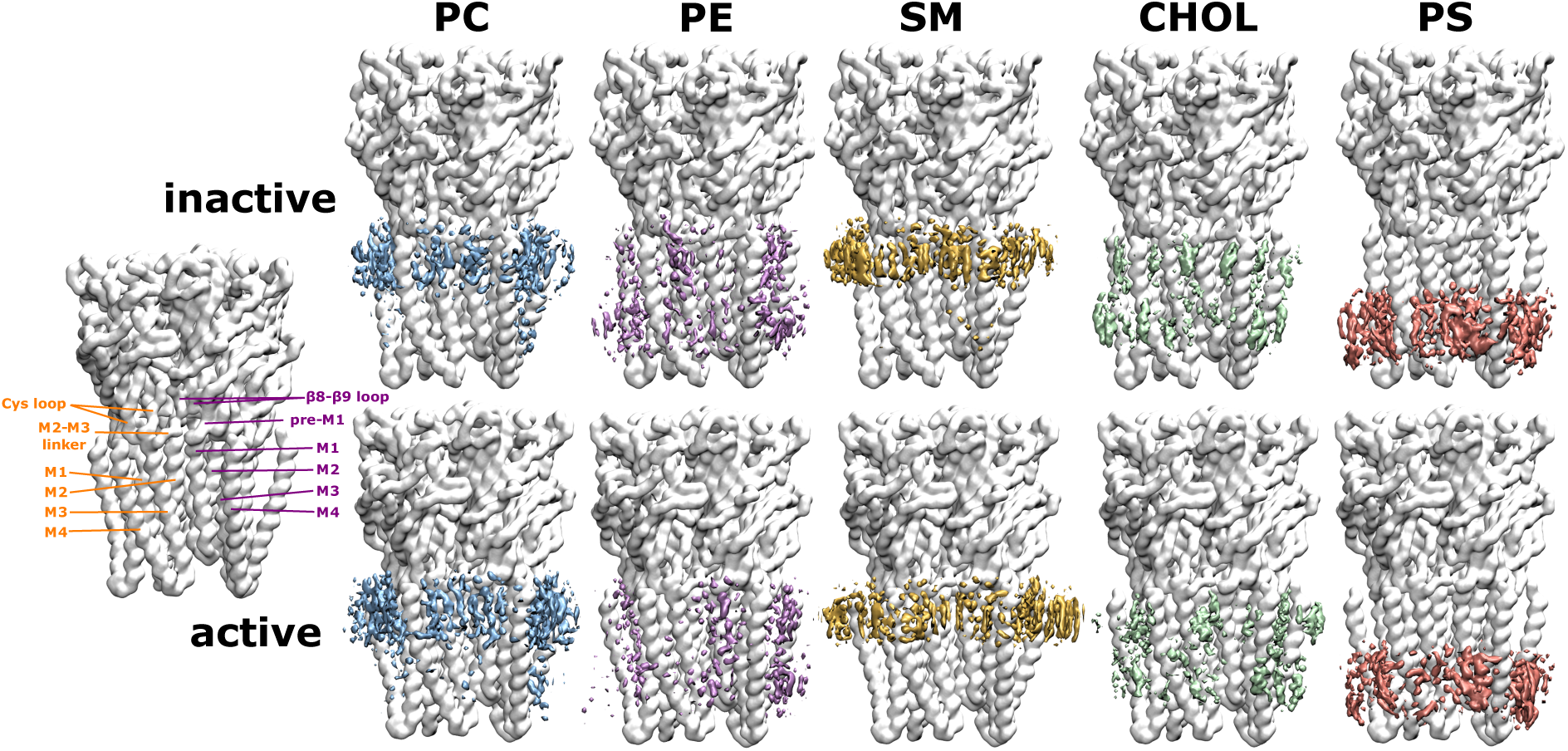
Densities of different lipid types in 3d space for the inactive and active state. For spatial orientation, the receptor structure on the left is labelled with key regions of adjacent subunits - principal (orange) and complementary subunit (purple). In order to better compare the individual lipid type differences for the two conformational states, the densities of each lipid type are shown at different isosurface values, such that they are reweighted as if the membrane consisted of equal amounts of each lipid type (the isosurface for the PC density is shown at a value of 6.3 (6.3) molecules/nm^3^, PE density at 5.4 (5.4) molecules/nm^3^, SM density at 3.7 (3.8) molecules/nm^3^, cholesterol density at 15 (15) molecules/nm^3^ and PS density at 5.8 (5.6) molecules/nm^3^ for the inactive state (active state)). In the active state, cholesterol can insert itself deep into a crevice that is opening up at the subunit interface between the (+)M3 and (-)M1 helices, where the binding site for the positive allosteric modulator ivermectin is located, indicating a possible positive modulatory role of cholesterol. The density of the anionic lipid type PS is higher in the inactive vs. the active state, suggesting that anionic lipids may aid in stabilizing the inactive receptor state. PC, PE and SM can all compete for the same regions on the receptor surface, which is not surprising given their similar chemical character.

In order to get a more detailed picture of the protein-lipid interactions of the first lipid shell, we calculated the lipid densities in 3-dimensional space (**Fig. 3)**. Striking state-dependent changes can be observed for cholesterol. Transitioning from the inactive to the active state allows cholesterol to penetrate into the subunit interface between the crevice of the (+)M3 and (-)M1 helices. Since this is where ivermectin, a known positive allosteric modulator, has been shown to bind, it suggests that cholesterol might act as an intrinsic membrane modulator for receptor activation. This is supported by experimental evidence that cholesterol is crucial for the activation of the nicotinic acetylcholine receptor (6, 12, 13, 15). Another functionally important lipid class for nicotinic acetylcholine receptors are anionic lipids. In our simulations, we have used PS, which is an important anionic lipid type in neuronal membranes that only occurs in the intracellular leaflet. We find that the PS density is higher in the inactive state compared to the active state, suggesting PS (and thus possibly in general anionic lipids) aid in stabilizing the inactive state over the active state. Overall, the different lipid types compete with each other for the same interaction regions on the receptor surface. Particularly, the neutral phospholipids PC, PE and SM can all interact with the same parts of the receptor. This is not surprising, as they are chemically more similar within themselves than compared to the steroid cholesterol or the anionic phospholipid PS (as discussed above).

### The duration of protein-lipid contacts is dependent on the receptor state

As a next step towards a more detailed understanding of the local strength of the protein-lipid interactions, we analysed the binding residues on the receptor for each specific lipid type. Binding residues would typically be identified as those residues with long contact times with lipid molecules. We therefore calculated the average duration of continuous contacts between a molecule of a given lipid species and each protein residue (using a 6 Å cut-off) as a proxy for local binding affinity (and mapped it onto the protein structure in **Fig. 4)**. Note that this can also be seen as the local residence time for a certain lipid molecule on a specific receptor residue.

**Figure 4.**
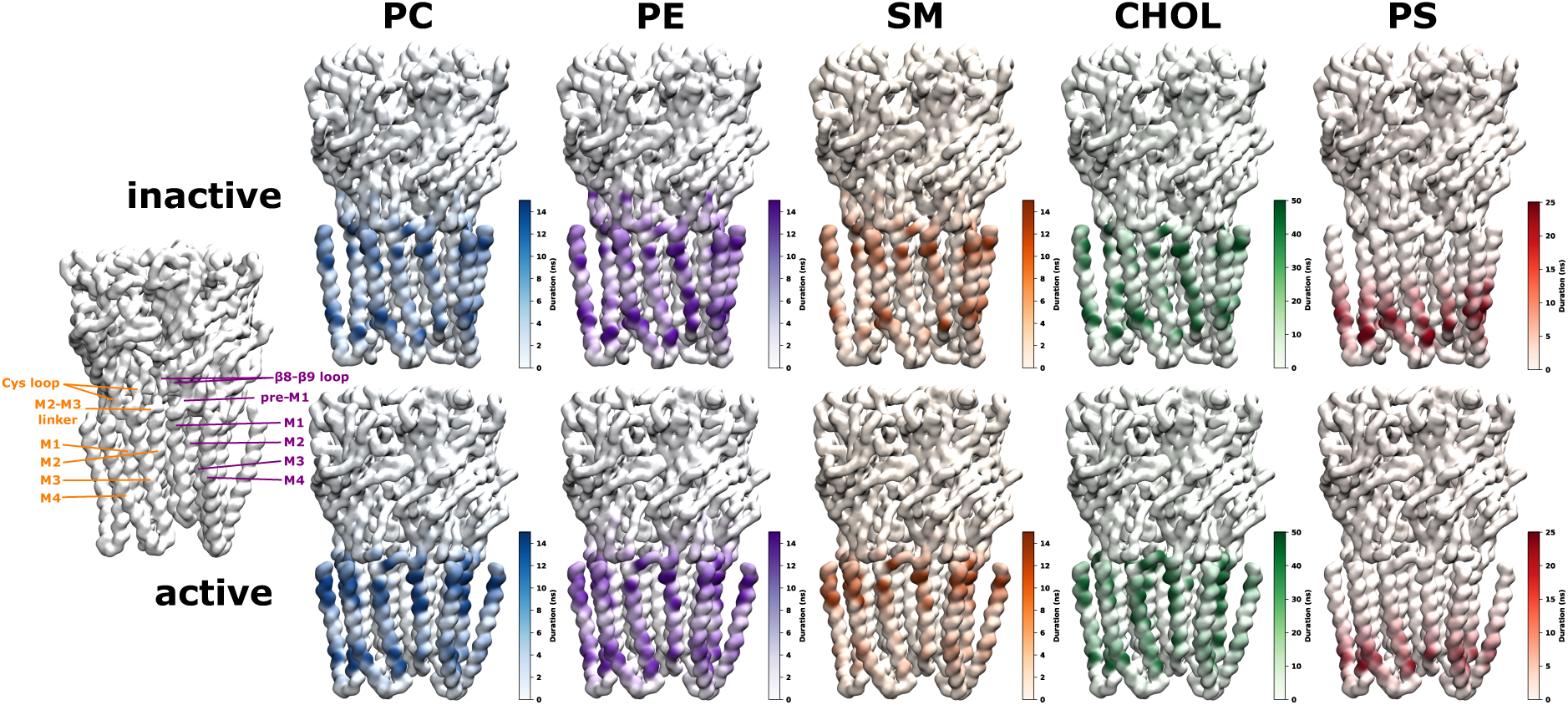
Mean duration of protein-lipid contact per residue for inactive and active state. Key regions of a principal (orange) and complementary subunit (purple) are shown on the left for spatial orientation. The colour scales were chosen so as to highlight the state-dependent differences per lipid type in an optimal way (the maximal contact duration can be longer than the maximum value on the colour scale, see **S1 Fig**). The per residue-interaction times are a proxy for the local binding affinity of a certain lipid type. Cholesterol forms the longest-lasting contacts with the receptor, followed by the anionic lipid PS, and then the chemically similar phospholipids PC, PE, SM, which supports the experimental evidence that cholesterol and anionic lipids are important for nAChR receptor function. Interestingly, all phospholipids interact with the ECD-TMD interface which is crucial for signal-transduction from the neurotransmitter binding ECD towards the TMD which contains the channel pore, thus, phospholipids could possibly modulate the conformational changes important for signal transduction at this pivotal interface.

Out of all lipid types, cholesterol engages in the longest contacts with the receptor (see also **S1 Fig**), indicating its most prevalent role as lipid interaction partner. The second most important lipid type in terms of interaction duration is PS. Finally, PC, PE and SM fall into a similar range of shorter contact times with the receptor. This hierarchy agrees with the experimental insight that cholesterol and a negatively charged lipid type (such as PS) are essential for nAChR function (45). As expected due to their chemical similarity, PC, PE and SM have very similar contact duration maps. For these lipids, we observe a trend towards longer interaction times in the extracellular half of the TMD when comparing the inactive with the active state. This suggests that the active state is stabilized by stronger interactions with these lipid types in the extracellular leaflet. Cholesterol forms much longer contacts in the active state at the subunit interface. In the active state, this site is more accessible and cholesterol forms longer contacts with the TMD regions forming this allosteric site, namely the extracellular halves of the (+)M3 helix, (+)M2 and (-)M1 helix as well as the (+)M2-M3 linker. Note that via direct interactions with the pore-lining M2 helix (which are also formed by the positive allosteric modulator ivermectin), cholesterol could directly influence the dynamics of the pore-lining M2 helices. Lastly, PS, which is only present in the lower leaflet, has longer contact times with the intracellular half of the TMD in the inactive vs. the active state.

### Membrane lipids can interact with the extracellular domain of the receptor

The PC, PE and SM interactions with regions of the ECD of the GlyR, are all with longer contact times to the inactive vs. the active state. The ECD-TMD junction of pLGICs is of particular importance for signal transduction, as it is the communication interface between the neurotransmitter binding ECD and the TMD containing the channel pore (see left panel of **Fig. 5**). Neurotransmitter binding is thought to trigger configurational changes in the β8-β9 loop (loop F), the β10 strand that precedes the M1 helix (pre-M1), the β1-β2 loop and the Cys loop (β6-β7 loop) in the ECD that form contacts with the M2-M3 linker of the TMD and thus transduce the signal into opening of the channel pore in the TMD. Note that loop C (β9-β10 loop) that covers the neurotransmitter binding site is known to close due to ligand binding and is directly linked to the β8-β9 loop and the pre-M1 region.

**Figure 5.**
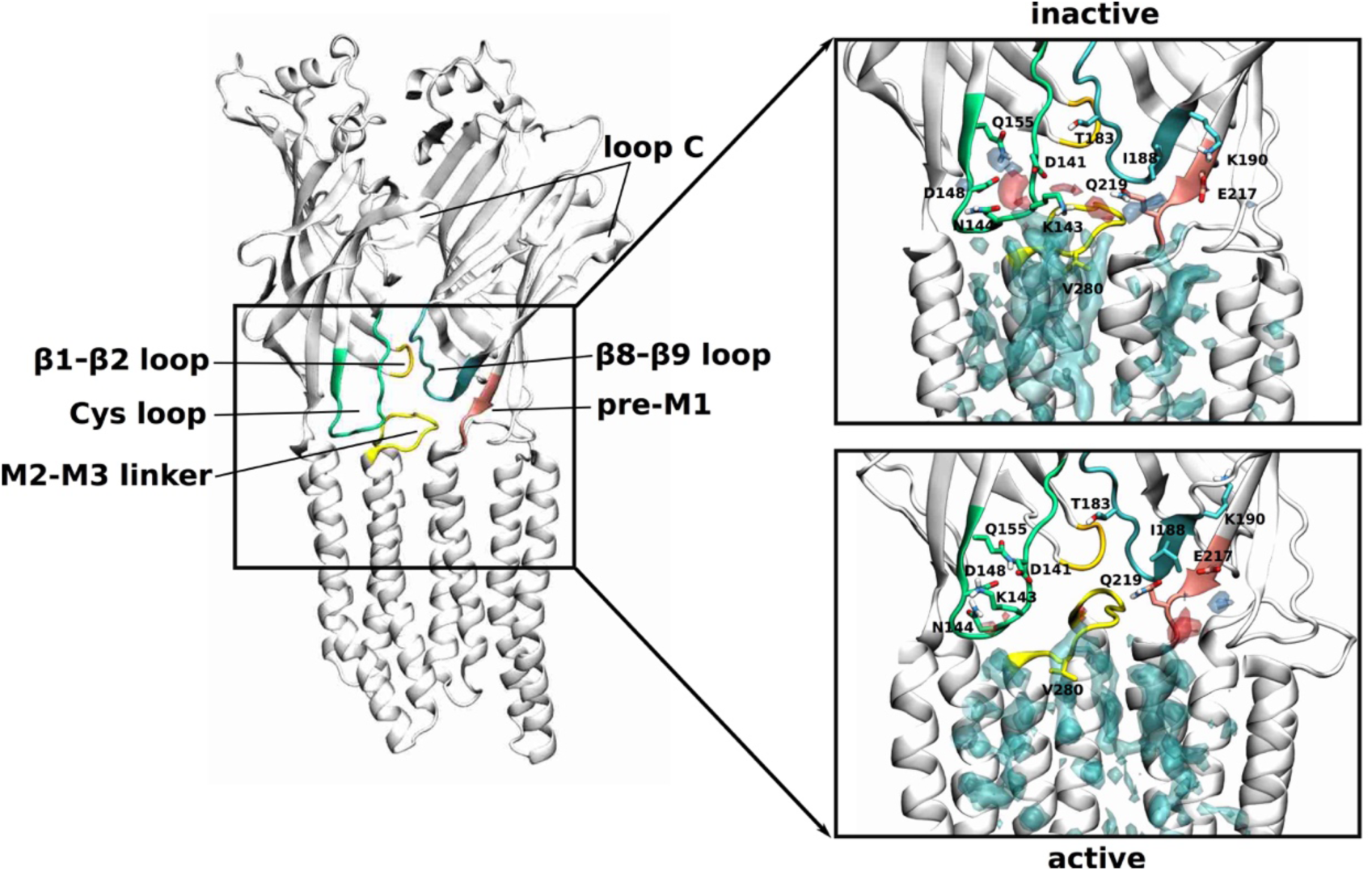
Phospholipid interactions with the ECD-TMD interface. Density arising from the positively charged headgroup beads (choline or ammonium group) is shown in transparent blue, from the negatively charged phosphate headgroup beads in transparent red and from the lipid tails transparent cyan. The densities corresponding to each lipid type are shown at different isosurface values, such that they are reweighted as if the upper membrane leaflet consisted of equal amounts of each lipid type (the isosurface for the PC density is shown at a value of 8.2 (8.2) molecules/nm^3^, PE density at 3.8 (3.6) molecules/nm^3^, SM density at 6.6 (6.8) molecules/nm^3^ for the inactive state (active state)). To show possible atomistic interactions, the initial model used as input to the coarse-grain simulations is shown here just as an indication. In the inactive state, the phospholipid headgroups interact with regions slightly further up towards the extracellular side of the ECD. These interactions are mediated mostly via polar side chains of the Cys loop, the β8-β9 loop as well as the part of the β10 strand that precedes the M1 helix (pre-M1). Note that the latter two are directly connected with loop C whose closing is key to neurotransmitter binding and thus, signal transduction towards the ECD-TMD interface.

Our simulations show that phospholipids (PC, PE and SM) interact with the receptor at precisely this crucial communication interface (**Fig. 5**). In the inactive state, the phospholipids preferentially interact with regions further up towards the extracellular side of the ECD. In particular, the Cys loop interacts with the zwitterionic head groups of phospholipids via the charged and polar residues D141, K143, N144, D148 and Q155. Another interaction hot spot is located more towards the complementary subunit, where the headgroups interact with Q219 and E217 from the pre-M1 region as well as T183 and K190 from the β8-β9 loop. The phospholipid densities are overall less far up towards the extracellular side of the ECD in the active compared to the inactive state. Moreover, the phospholipid head group densities are less defined near the Cys loop, indicating more flexibility as well as a weaker interaction strength. In the active state, the phospholipid head group density is much more defined near Q219 and E217 of the pre-M1 region of the complementary subunit, indicating more stable interactions at this site. Furthermore, a more defined lipid binding site appears in the active state at the subunit interface that is not as clearly separated in the inactive state. This density originates mostly from SM (compare also additional density spots of SM at the subunit interfaces in the extracellular leaflet in the active state in **Fig. 2**) and could be due to favourable interactions between SM and CHOL. In summary, the phospholipid density analysis at the ECD-TMD interface suggests that in the inactive state, the phospholipid head groups form more stable interactions with the Cys loop but in the active state they form more stable interactions with the pre-M1 region.

Although for the GlyR, no phospholipid density has been fully resolved in the structures available at present, structural data of other members of pLGIC superfamily with phospholipid or detergent sites at the ECD-TMD interface does exist and a comparison is made in the SI (**S1 Text, S2 Fig**).

### Cholesterol can bind to the binding site of the positive allosteric modulator ivermectin

Cholesterol has the longest-lived interactions and the binding site corresponds closely to the binding site of ivermectin, which is between the principal and complementary subunit via the (+)M3 and (-)M1 helices and forms important contacts with the pore lining (+)M2 helix and the (+) M2-M3 linker (Fig. 6). This site is not accessible in the inactive receptor state, but opens up in the active or desensitized states of the receptor. While cholesterol preferentially interacts at the mouth of the ivermectin binding site in the inactive state, it only deeply penetrates it in the active state, as can be seen in **Fig. 2-4**. In the active state, cholesterol density overlaps with the position of structurally resolved ivermectin and the density we observe in our simulations can be assigned to 2 cholesterol molecules. Residues previously identified to be involved with ivermectin binding, such as A288 from the (+)M3 helix, I225, I229, P230 and L233 from the (-)M1 helix as well as V280 from the (+)M2-M3 also interact with the density arising from the hydrophobic cholesterol steroid skeleton (**Fig. 6**). The cholesterol density overlays well with the spiroketal and disaccharide part of ivermectin. The density suggests that one cholesterol molecule forms polar interactions via its hydroxy group with D284 from the (+)M3 helix, while a second cholesterol molecule can form polar actions via its hydroxy group with polar backbone atoms of the (+)M2-M3 linker, namely the backbone amine group of V280, the carbonyl group of S278 and the carbonyl group of A277 (Fig. 6). Interestingly, the polar atoms of ivermectin’s disaccharide tail are also orientated towards this region of the (+)M2-M3 linker. The depth of penetration at this site by modulatory molecules is interesting and may be related to their potency as it is also known that ivermectin has higher potency at GluCl, where it is buried ca. 1 Å deeper in the cleft than in the human α3 GlyR. Atomistic MD simulations have previously also identified preferred cholesterol binding to the ivermectin binding site in the GABA_A_R (41). However, on a sub-microsecond time scale, the sampling of the highly dynamic interactions was much more limited.

**Figure 6.**
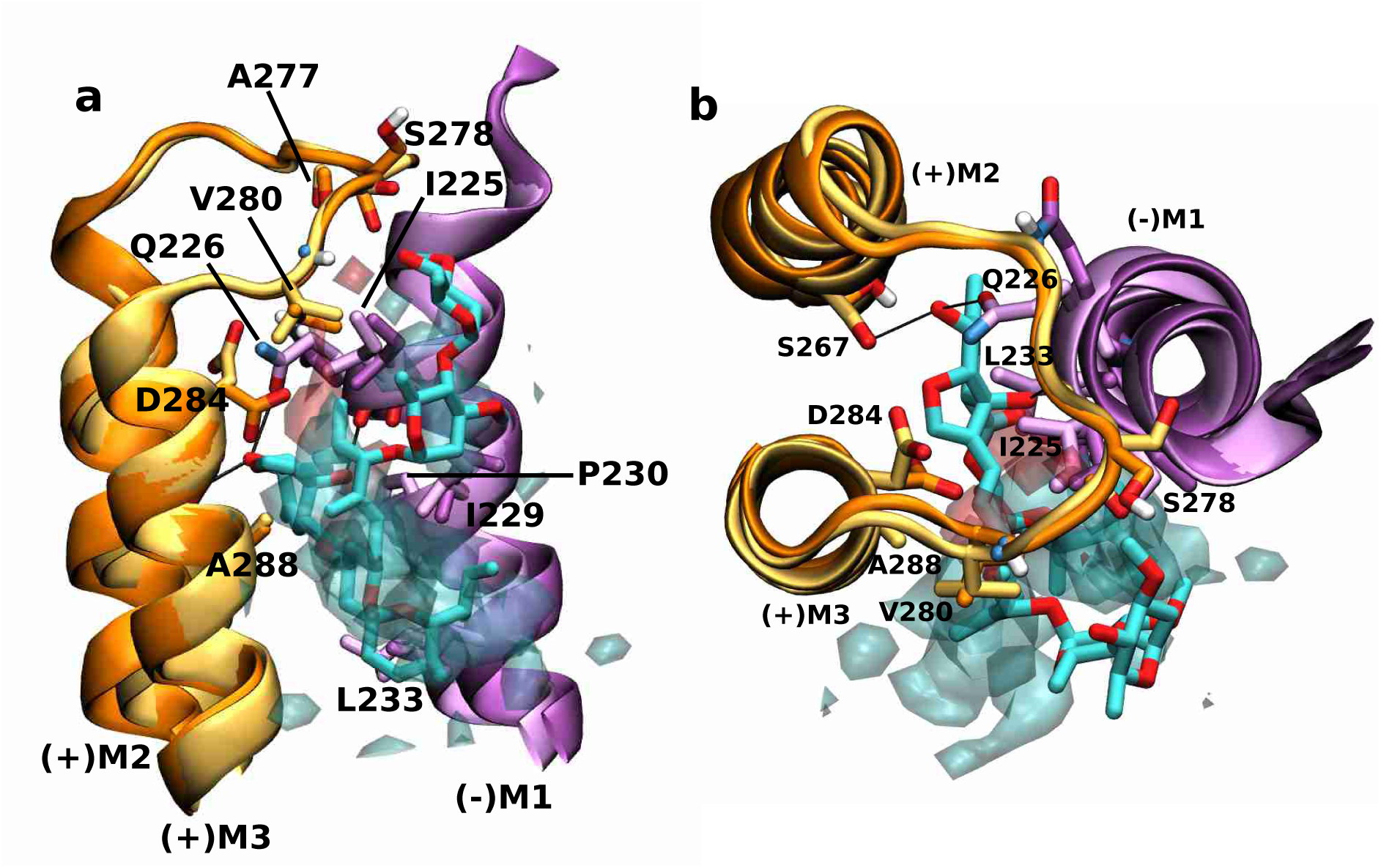
Overlay of cholesterol density with positive allosteric modulator ivermectin at the subunit interface. Overlay of the human α1 GlyR model (darker colours) and cholesterol density (transparent, hydroxy group density in red, remaining hydrophobic steroid skeleton in cyan) from simulations in the active state with the currently highest resolution crystal structure of human α3 GlyR (PDB 5VDH) (lighter colours) with ivermectin bound. Shown is the ivermectin binding site at the interface of adjacent subunits with the (+)M2 and (+)M3 helices of the principal subunit (orange) and the (-)M1 helix of the complementary subunit (purple) forming a crevice and the (+)M2-M3 linker acting as a lid towards the extracellular side. Ivermectin and important residues for ivermectin or cholesterol binding are shown explicitly and key interactions are highlighted by black lines. **a** Side view almost in parallel to the membrane and **b** view from the extracellular side towards the intracellular side.

### The binding mode of the anionic lipid PS in GlyR differs from the PIP2 binding mode in the GABA_A_ receptor

Comparison of the PIP2 binding site at one of the α1 subunits of the human α1β3γ2 GABA_A_ receptor (PDB 6I53) overlaid with the density originating from the anionic lipid PS from our active state simulations reveals the overall location of the anionic binding site is similar at the intracellular end of the M4 and M3 helices (**Fig. 7**). However, the binding modes differ between PIP2 and PS. The most striking difference in the receptor structures is that the M4 helix of the GlyR structure maintains its helical secondary structure for 2 more turns towards the intracellular side than the M4 helix of the GABA_A_R structure. As a consequence, this region sterically prohibits access of PS in the GlyR, which is precisely where the inositol-1,4,5-trisphosphate (InsP3) head group of PIP2 interacts extensively with the protein backbone of the GABA_A_R. Moreover, the positively charged side chains of GABA_A_R (R249, K312, R313) that electrostatically stabilize the negatively charged InsP3 head group of PIP2 in this region are either not present in human α1 GlyR or in orientations too far away to engage in such interactions (R249 and R309). In GlyR, the negatively charged phosphatidylserine headgroup of PS forms stable electrostatic interactions slightly further towards the extracellular side of the M4 helix with the positively charged side chains R391 and K394. Furthermore, the InsP3 headgroup of PIP2 is overall larger and carries a two-fold negative charged, whereas the phosphoserine headgroup of PS is smaller and carries only a single charge, which is also likely to have an effect on the different binding locations for these headgroups.

**Figure 7.**
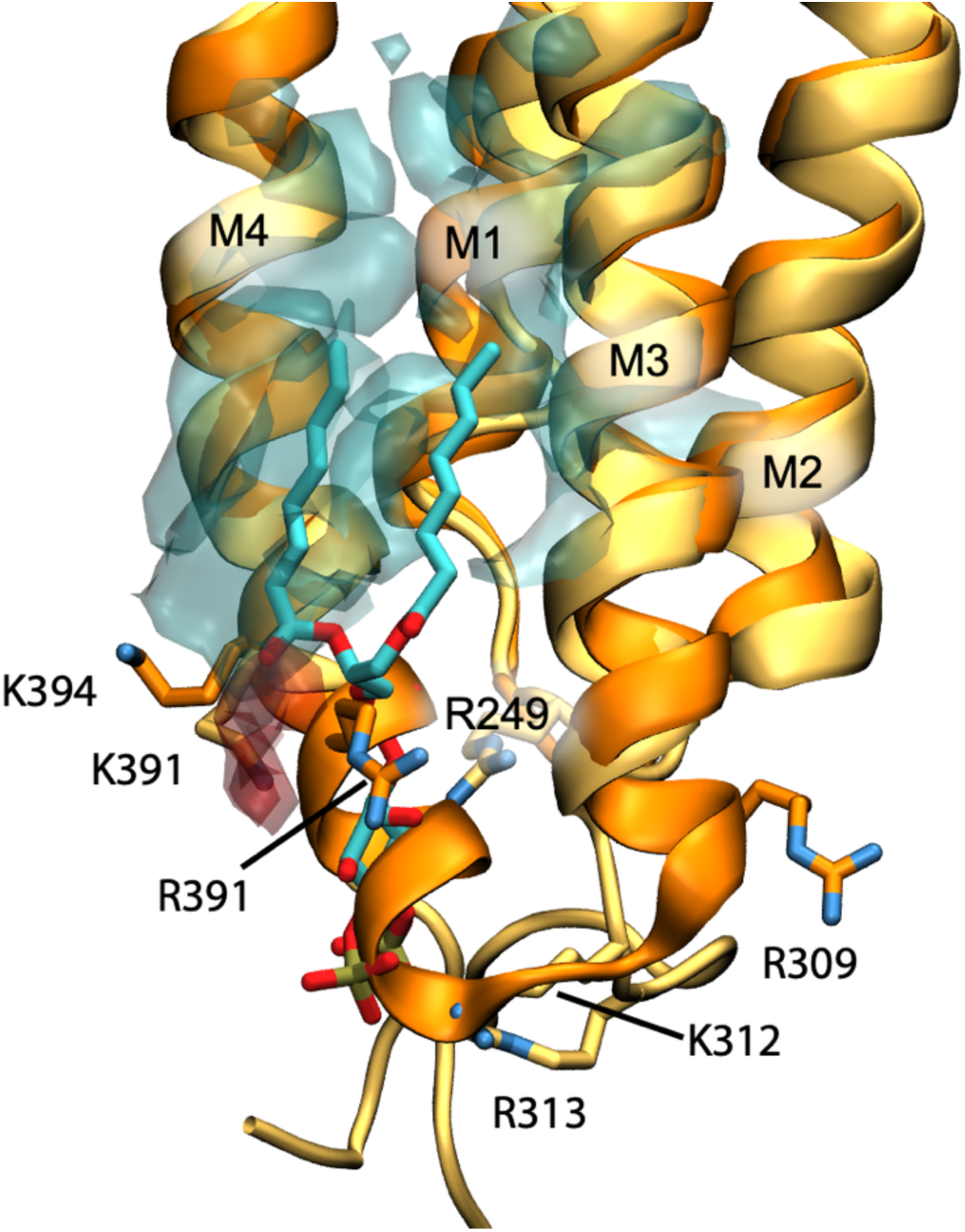
Overlay of PS density with PIP2 near the M4 helix. Overlay of the human α1 GlyR model (darker colours) and PS density (transparent, negatively charged head group density in red, lipid tail density in cyan) from simulations in the active state with the cryo-EM structure of human α1β3γ2 GABA_A_ receptor (PDB 6I53) (lighter colours) with PIP2 bound. Shown is the PIP2 binding site at one of the α1 subunits. PIP2 and important residues for anionic lipid binding are shown explicitly.

## Discussion

We have computationally reconstituted the GlyR in a realistic model of a neuronal membrane and performed CG-MD simulations to probe and characterize the molecular level detail of the protein-lipid interactions in two key functional receptor states (inactive and active). Our simulations show that lipid interactions with GlyR are dependent on the receptor state. The differences in the residence times between the inactive and active state are a proxy for changing lipid binding affinities at their respective interaction sites that must occur during receptor activation. This suggests that the lipid environment itself might play an active role in regulating the activation of the receptor. Of particular note, we characterized so far unreported protein-lipid interactions at the ECD-TMD interface, suggesting that lipids might play a modulatory role at this communication interface in the activation process of the receptor. Two phospholipid interaction sites at the ECD-TMD interface coincide well with known phospholipid or detergent binding sites in GLIC and GABA_A_R and, in addition, we predict a third phospholipid binding site at the extracellular half the TMD at the subunit interface in the active state of GlyR. Furthermore, our simulations show that cholesterol binds to the binding site of the positive allosteric modulator ivermectin, indicating that cholesterol might also have a positive modulatory function. Finally, we identify a binding site for the anionic lipid PS at the intracellular leaflet half of the TMD on the surface of the M4 and M3 helices which roughly coincides with the location of PIP2 binding in human GABA_A_R. This region was also identified as important for the binding of PE to ELIC (17).

The nAChR is the best studied member of the pLGIC superfamily with regards to protein-lipid interactions (46). Cholesterol and anionic lipids have both been shown to be essential for nAChR function (12-15). NAChRs reconstituted in pure PC or PE membranes bind agonist, but do not open. The unresponsive state of nAChR has been called the uncoupled state. However, mechanistic insight into the molecular details that characterize the uncoupled vs. the responsive (or coupled) state has been elusive so far. Lipid dependency seems to be an important theme for the whole pLGIC superfamily, as the function of GABA_A_Rs is also dependent on cholesterol levels (47).

Since current evidence suggests that lipids can modulate nAChR function via interactions with the M4 helix as well as the adjacent M1 and M3 helices, the so called M4 lipid-sensor model has emerged (21). It hypothesizes that the uncoupled state is structurally characterized by the M4 helix being more distant to its neighbouring M1 and M3 helices and thus the extracellular post-M4 cannot form interactions with the Cys loop of the ECD. Lipids could influence the binding of the M4 helix to the adjacent M1/M3 surface and thus either favour the coupled or uncoupled state (48). As aromatic residues lining the contact interface of the M4 and M1/M3 helices have been shown to be the main driving force of M4 helix binding to the M1/M3 surface during the folding of GlyR (22), it has further been hypothesized that the varying number of these π-π interactions in pLGIC members is linked to the degree of dependence on the lipid environment (21). The more aromatic residues support the M4-M1/M3 binding, the less dependent is a receptor on its lipid environment. This hypothesis was given further support recently by elegant work on the ELIC channel which revealed a lipid binding site in this region, which can be linked to the intrinsic flexibility of the M4 helix (17).

**Fig. 8** gives an overview of the varying degree of aromatic residues lining the M4-M1/M3 interface across representatives of the pLGIC superfamily. In the human α1 GlyR aromatic residues line the M4-M1/M3 interface (**Fig. 8a**) in an evenly distributed manner, leading to a tight binding of the M4 helix to the M1/M3 interface and thus, according to the M4 lipid-sensor model, suggesting little dependence on the lipid environment. A similarly aromatic rich interface is found in the β3 GABA_A_R (**Fig. 8b**). The prokaryotic channel GLIC has only 8 aromatic residues at the M4-M1/M3 interface with a striking gap at the extracellular half of the TMD (**Fig. 8c**), indicating weaker binding of the upper half of the M4 helix. On the other end of the spectrum are the nAChR (α4 subunit shown in (**Fig. 8d**)) and ELIC (**Fig. 8e**) with only 6 and 4 aromatic residues, respectively, suggesting a weak binding of the M4 helix and the M4 lipid-sensor model would predict a strong dependence on the lipid environment. Indeed, nAChR and ELIC have been demonstrated to be highly dependent on their lipid environment and do not undergo agonist-induced opening in minimal PC membranes, whereas GLIC remains functional (21). While experimental data for GlyR and GABA_A_R in this respect are not available, their larger number of aromatic contacts along the M4-M1/M3 interface than GLIC suggest that they would remain functional in minimal PC membranes.

**Figure 8.**
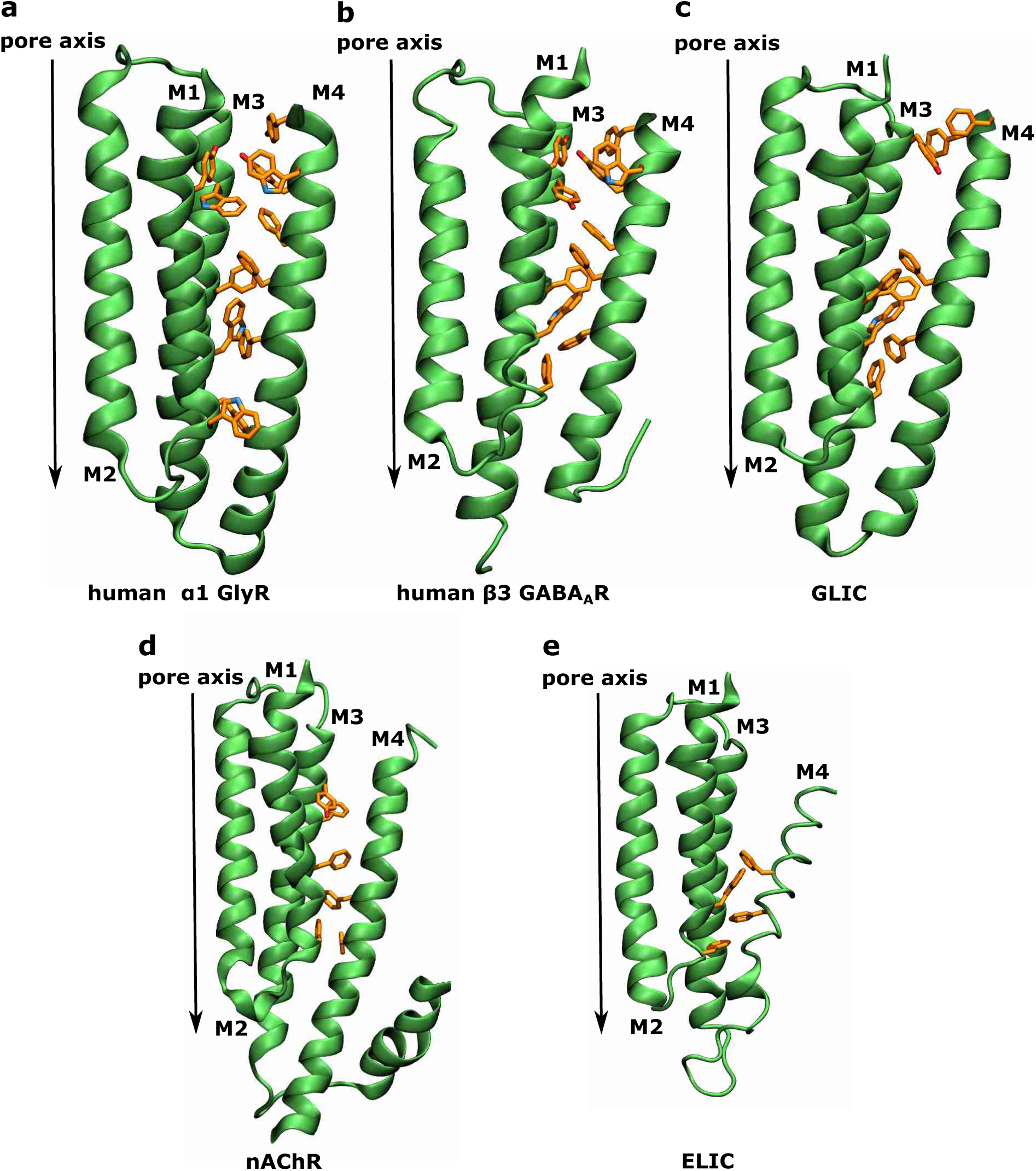
Aromatic residues at the M4-M1/M3 interface of different pLGICs. TMD region of a single subunit viewed in parallel to thse membrane plane with aromatic residues at the M4-M1/M3 interface explicitly shown in orange for **a** human α1 GlyR (human homology model from PDB 3JAE (28)), **b** human β3 GABA_A_ receptor (from PDB 6I53 (23)), **c** GLIC (from PDB 6HZW (25)), **d** α4 human nicotinic acetylcholine receptor (from PDB 5KXI (49)) and **e** ELIC (from PDB 2VL0 (50)).

Interestingly, the analysis of aromatic contacts at the M4-M1/M3 interface explains our observation that phospholipids bind to the membrane facing surface of the M1 and M4 helices of GlyR rather than inserting in between the M4-M1/M3 interface. The plethora of aromatic contacts between the M4 helix and the M1/M3 surface binds the M4 helix strongly to the receptor in GlyR. This may also be the reason why similar surface binding of phospholipids is observed in the aromatic-rich GABA_A_R.

The protein-lipid interactions described here form the basis for further experimental studies to address their physiological functional consequences *in vivo*. From a pharmacological perspective, the detailed picture of protein-lipid interactions for the GlyR as a pLGIC representative forms the basis to guide future design principles for lipid-like drugs that can partition into the membrane and bind to allosteric sites within the TMD or even at the ECD-TMD interface.

With the ongoing cryo-EM structural revolution, our structural knowledge of interacting lipids with pLGICs will undoubtedly increase with more high resolution structures being resolved that show electron density for lipid molecules. Since it is still difficult to identify the exact lipid type from the electron density data alone, MD simulations will be an important complementary technique in order to both identify the molecular identities of the lipid electron density as well as provide insight into the dynamic lipid interactions with the receptor (10).

## Methods

### Homology Modelling

The model of the active conformation of human α1 GlyR was constructed from the structure of *Danio rerio* α1 GlyR (glycine-bound open state; PDB: 3JAE (28)) as previously reported (51). The model in the inactive conformation was based on the *human* α3 GlyR (strychnine-bound closed state; PDB: 5CFB). Sequences were aligned using Muscle (52) and adjusted manually. 100 models were generated via MODELLER version 9.16 (53) and from the intersection of the top 10 of both molecular PDF and DOPE scores (54), the model with the best QMEAN score (55) was chosen for simulation. The pdb2gmx tool of Gromacs 5.1 (56) was used to add hydrogens with standard protonation states at pH = 7.4.

### System Set Up

The homology models of GlyR were converted to coarse-grained MARTINI representations via martinize (34). Using the insane script (57), each of the coarse-grained protein structures was separately embedded in an asymmetric neuronal model membrane using a hexagonal prism as the periodic simulation box, solvated in water and sodium and chloride ions were added to neutralise the net charge and simulate a physiological concentration of 150 mM (see **Fig. 1**). The membrane model is based on (43) and was simplified to consist of representatives of the five most essential lipid species as specified in **Table 1**. For each receptor state, 10 systems were set up with the same number of molecules per species, but with different random seeding of the lipid positions in the membrane plane to improve sampling. No constraints were imposed to maintain the asymmetric lipid distribution across leaflets, but based on previous experience only cholesterol can interchange the leaflets on timescales of tens of μs.

**Table 1:**
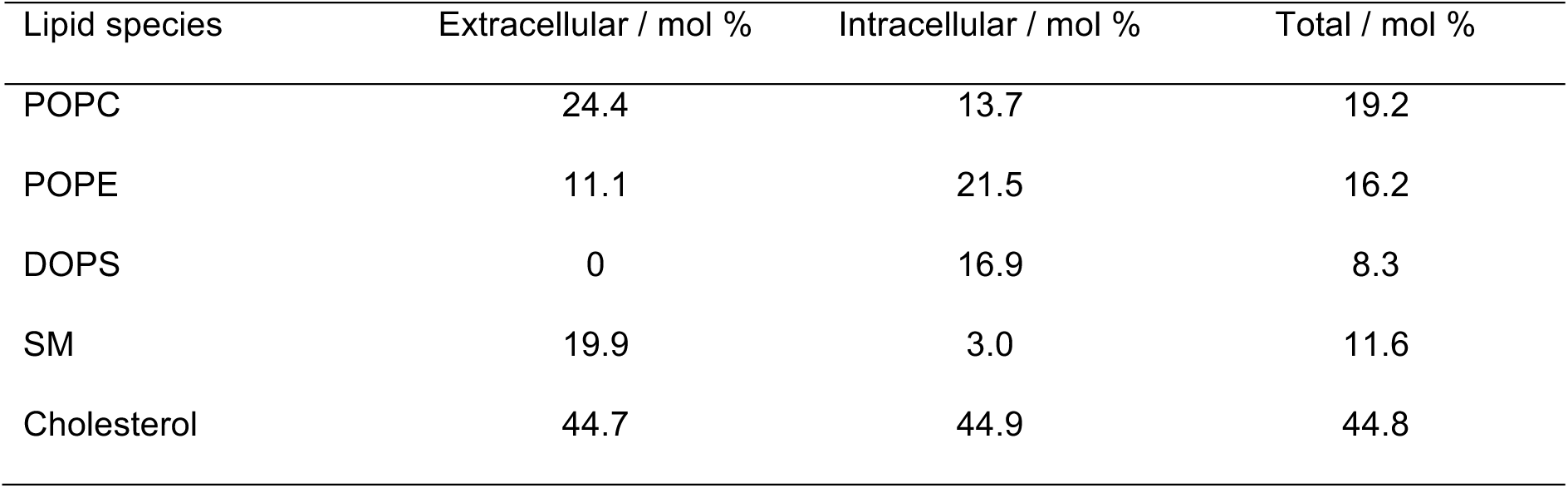
Composition of a human neuronal model membrane (simplified to 5 essential lipid types based on the model by Ingolfsson *et al* (43)).

| Lipid species | Extracellular / mol % | Intracellular / mol % | Total / mol % |
| --- | --- | --- | --- |
| POPC | 24.4 | 13.7 | 19.2 |
| POPE | 11.1 | 21.5 | 16.2 |
| DOPS | 0 | 16.9 | 8.3 |
| SM | 19.9 | 3.0 | 11.6 |
| Cholesterol | 44.7 | 44.9 | 44.8 |

### Coarse Grained Molecular Dynamics (CG-MD) Simulations

All CG-MD simulations were performed with the Martini force field version 2.2 for protein and version 2.0 for lipids (58). Gromacs 2018 (56) was used as molecular dynamics engine for all simulations and periodic boundary conditions were applied in all three spatial dimensions. After energy minimisation, the systems were simulated at 310 K and 1 bar for 40 μs. Simulation details were based on the recommended new-rf.mdp parameters (34) and appropriately adjusted. Harmonic restraints with a force constant of 1,000 kJmol^-1^nm^-2^ were placed on all protein particles to preserve the conformational state close to the original crystal or cryo-EM structure.

### Analysis and Visualization

Trajectory analysis and probability density calculations were performed on the last 20 μs out of a total of 40 μs simulation time using frames at a separation of dt = 1 ns and averaged over all 10 repeats per conformational state. The contact-duration data was additionally averaged over all five subunits. A cut-off of 6 Å (corresponding to the first lipid shell) was used to define a protein-lipid contact. Software used for analysis was MDAnalysis (59), MDTraj (60) and in-house scripts, part of which are now publicly available on https://github.com/wlsong/PyLipID. Visualization was performed in VMD (61).

## Acknowledgements

We thank our colleagues Wanling Song and Anna Duncan for useful discussions. MAD received funding by the Oxford Wolfson Marriott Graduate Scholarship in Biochemistry, the Studienstiftung des deutschen Volkes, the Goodger and Schorstein Research Scholarship in the Medical Sciences as well as the Vice-Chancellors’ fund of the University of Oxford. This project made use of computation time on JADE (EP/P020275/1) via HECBioSim (http://www.hecbiosim.ac.uk), supported by EPSRC (grant no. EP/R029407/1). PCB thanks the MRC for funding (BB/S001247/1).

## Author Contributions

MAD and PCB conceived the study. MAD performed and analysed the MD simulations. MAD and PCB interpreted the results and wrote the paper.

## Supporting Information

**S1 Fig. Mean duration of protein-lipid contact per residue for inactive and active state**. Averaged over last 20 μs (out of 40μs) of 10 repeats per conformational state and over the 5 subunits. A cut-off of 6 Å is used, which corresponds to the first lipid shell.

**S1 Text. Comparison of lipid binding sites to other pLGIC members**.

**S2 Fig. Overlay of phospholipid densities at the ECD-TMD interface with experimental structures**. Overlays of the human α1 GlyR model (darker colours) and phospholipid density (transparent, phosphate headgroup density in red, choline/ammonium headgroup density in blue, tail density in cyan) from simulations in the active state with experimentally resolved structures of the pLGIC superfamily (lighter colours) with phospholipid or detergent bound. Three binding sites are observed in simulations (labelled 1, 2 and 3). Lipid binding has been observed in the corresponding regions of sites 1 and 2 for other members of the pLGIC family, but site 3 may be specific to the GlyRs a Overlay with GLIC crystal structure (PDB 6HZW) which has a detergent molecule bound that interacts with its polar headgroup with the Cys loop, while its hydrophobic tail mainly interacts with the M3 helix. b Overlay with GLIC crystal structure (PDB 6HZW) which has a phosphocholine lipid bound that interacts with its polar headgroup with the pre-M1 region as well as the Cys loop, while its hydrophobic tails bridge the interface between the M4 and M1/M3 helices. c Overlay with the α1 subunit of the human GABA_A_ receptor cryo-EM structure (PDB 6I53), which has a phospholipid bound that interacts with its polar headgroup with the pre-M1 region, while its hydrophobic tails interact mainly with the M1 helix. d Overlay with the β3 subunit of the human GABA_A_ receptor cryo-EM structure (PDB 6I53), which has a phospholipid bound that interacts with its polar headgroup with the pre-M1 region, while its hydrophobic tails interact with both the M1 and M4 helices.

**S1 Movie. Side view of rock and roll movie of cholesterol overlaid on ivermectin binding site**.

**S2 Movie. Top view of rock and roll movie of cholesterol overlaid on ivermectin binding site**.

**S1 README. Text file describing input files and trajectory files to be deposited on zenodo**.**org**.

